# Elucidating the CodY regulon in *Staphylococcus aureus* USA300 substrains

**DOI:** 10.1101/2021.01.08.426013

**Authors:** Ye Gao, Saugat Poudel, Yara Seif, Zeyang Shen, Bernhard O. Palsson

## Abstract

CodY is a conserved broad acting transcription factor that regulates the expression of genes related to amino acid metabolism and virulence in methicillin-resistant *Staphylococcus aureus* (MRSA). CodY target genes have been studied by using *in vitro* DNA affinity purification and deep sequencing (IDAP-Seq). Here we performed the first *in vivo* determination of CodY target genes using a novel CodY monoclonal antibody in established ChIP-exo protocols. Our results showed, 1) the same 135 CodY promoter binding sites regulating 165 target genes identified in two closely related virulent *S. aureus* USA300 TCH1516 and LAC strains; 2) The differential binding intensity for the same target genes under the same conditions was due to sequence differences in the same CodY binding site in the two strains; 3) Based on transcriptomic data, a CodY regulon comprising 72 target genes that are differentially regulated relative to a CodY deletion strain, representing genes that are mainly involved in amino acid transport and metabolism, inorganic ion transport and metabolism, transcription and translation, and virulence; and 4) CodY systematically regulated central metabolic flux to generate branched-chain amino acids (BCAAs) by mapping the CodY regulon onto a genome-scale metabolic model of *S. aureus*. Our study performed the first system-level analysis of CodY in two closely related USA300 TCH1516 and LAC strains giving new insights into the similarities and differences of CodY regulatory roles between the closely related strains.

**Importance:** With the increasing availability of whole genome sequences for many strains within the same pathogenic species, a comparative analysis of key regulators is needed to understand how the different strains uniquely coordinate metabolism and expression of virulence. To successfully infect the human host, *Staphylococcus aureus* USA300 relies on the transcription factor CodY to reorganize metabolism and express virulence factors. While CodY is a known key transcription factor, its target genes are not characterized on a genome-wide basis. We performed a comparative analysis to describe the transcriptional regulation of CodY between two dominant USA300 strains. This study motivates the characterization of common pathogenic strains and an evaluation of the possibility of developing specialized treatments for major strains circulating in the population.

## Introduction

*Staphylococcus aureus* is a ubiquitous, Gram-positive pathogen that causes a diverse range of bacterial infections, from skin and soft tissue infections to potentially fatal infections, such as pneumonia, endocarditis, osteomyelitis, sepsis, and toxic shock syndrome (1). Coupled with the growing prevalence of methicillin-resistant *S. aureus* (MRSA) strains, the worldwide threat posed by this pathogen is obvious (2). Current research provides insights into important features of these strains, including antibiotic resistance and extensive virulence factors. So far, a number of known virulence factors consist of surface-associated proteins, such as microbial surface components, and secreted proteins, like hemolysins, immunomodulators, and many exoenzymes (3). To combat the worldwide spread of *S. aureus*, significant effort is being focused on the investigation of the transcription factors that control virulence factors during infection (4, 5). With different strains circulating in the population, it is important to understand the fundamental differences between them.

CodY is an important broad acting transcription factor in *S. aureus* (6, 7). While the primary known role of CodY is to regulate the metabolic genes in response to cellular branched chain amino acid and GTP concentrations, it also controls the expression of several virulence factors, acting as a bridge between metabolism and virulence (6, 8–11). The CodY regulon, estimated to consist of 150 to more than 200 genes, can vary in size between *S. aureus* strains, and CodY can even have opposite effects on the expression of the same gene in different strains (10, 12, 13). At present, direct determination of CodY DNA-binding sites using ChIP-exo and the identification of the corresponding target genes has not been achieved.

To address this challenge, we developed a reliable ChIP-grade monoclonal antibody and conducted in *vivo* genome-wide experiments (ChIP-exo) to identify 165 identical CodY target genes in two common *S. aureus* USA300 isolates (TCH1516 and LAC). To reconstruct the CodY regulon, we compared RNA-seq profiles of the wild-type strain and *codY* mutant. To examine the network level effects of CodY, we used a genome-scale metabolic model to simulate the flux state of central carbon metabolism, demonstrating the regulatory activities of CodY to generate BCAA. Our study used a comprehensive pipeline to characterize the CodY regulon between closely related USA300 substrains that included genetic parameters and network level computational models.

## Results

### Comparative genomic analysis of *S. aureus* USA300 reveals a high level of identity between two closely related strains, TCH1516 and LAC

The community-associated methicillin-resistant *S. aureus* (CA-MRSA) clones belonging to the USA300 lineage have become the dominant sources of MRSA in the United States (14). They are distinct from other *S. aureus* clones such as USA200 (UAMS-1), USA400 (MW2), and ST8 MSSA (Newman) (14). Two dominant CA-MRSA USA300 strains, TCH1516 and LAC, isolated in Los Angeles and Houston, respectively, were chosen to study CA-MRSA clones as they represent well characterized strains in the USA300 lineage (15). These strains have become the epidemic clones spreading in the community (16).

First, using whole-genome alignment, we characterized the genomes of TCH1516 and LAC, with lengths of ∼2.872 Mb and ∼2.878 Mb, respectively (**Table 1**). We found a high level of identity at the nucleotide level between these two closely related strains, and on average, they displayed 99.5% identity for all coding genes (**Figure 1A**). Further, MUMmer was used to check for the synteny between the strains. We produced a MUMmer dot plot resulting from the alignment of their chromosomal sequences, demonstrating that they have high similarity between the genomes and large scale inversions located at positions ∼ 2.18 Mb (TCH1516) and ∼ 2.20 Mb (LAC) (**Figure 1B**). Overall, these data revealed that they have a high degree of sequence similarity throughout their genome sequences.

**Table 1.** Comparison of annotation features in *S. aureus* USA300 lineage

|  | *S. aureus* TCH1516 | *S. aureus* LAC |
| --- | --- | --- |
| Total sequence length (bp) | 2,872,915 | 2,878,171 |
| Total chromosome and plasmids | 3,<br>NC_010079.1 (chromosome)<br>NC_012417.1 (plasmid 1)<br>NC_010063.1 (plasmid 2) | 2,<br>CP035369.1 (chromosome)<br>CP035370.1 (plasmid) |
| Gene (total) | 2920 | 2892 |
| CDS (total) | 2841 | 2802 |
| Genes/CDS (coding) | 2763 | 2733 |
| RNA | 79 | 90 |
| rRNAs (5S, 16S, 23S) | 6, 5, 5 | 7, 6, 6 |
| tRNAs | 59 | 67 |
| ncRNAs | 4 | 4 |
| Pseudo gene (total) | 78 | 69 |

**Figure 1.**
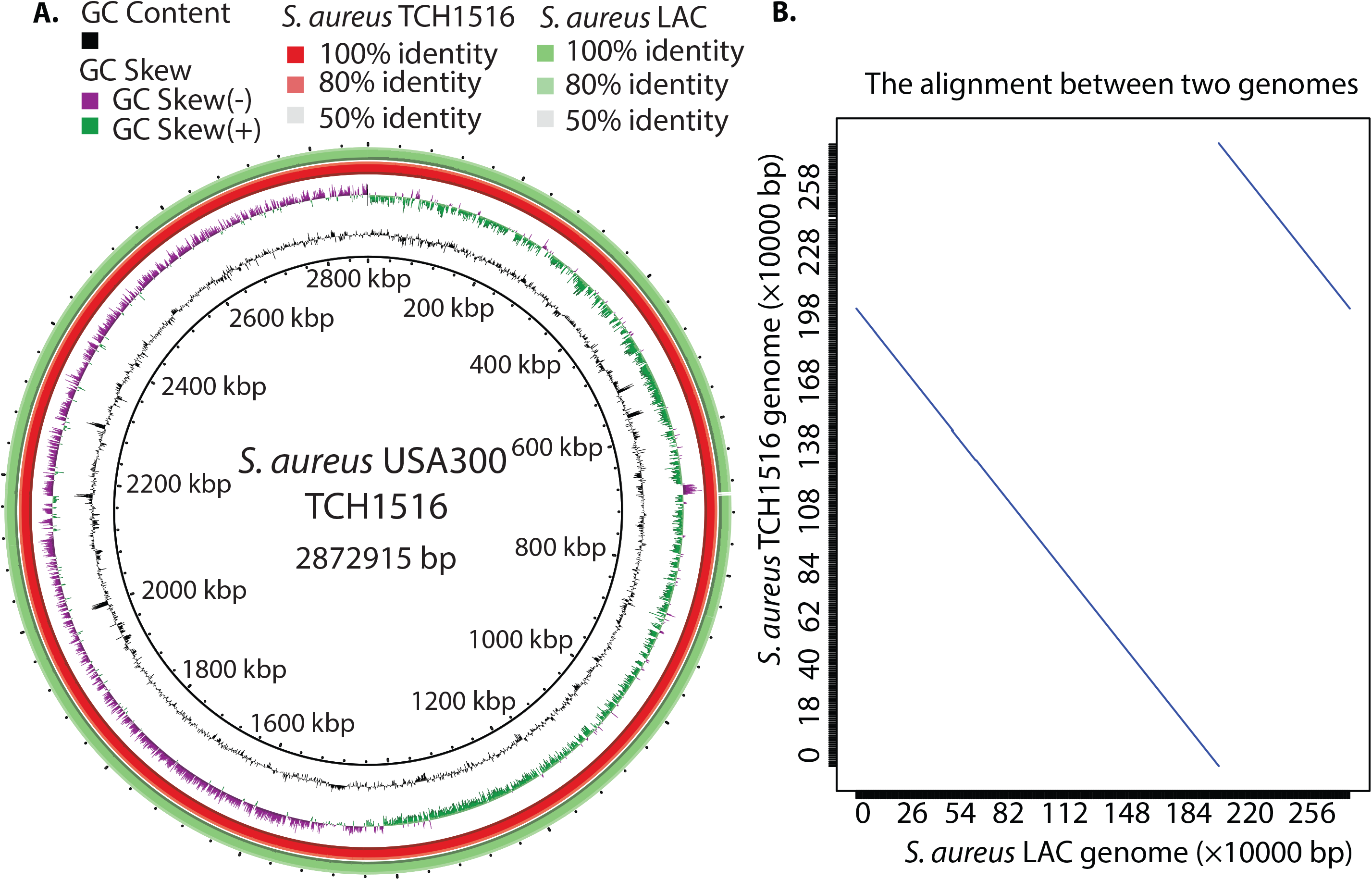
The comparison of two dominant *Staphylococcus aureus* USA300 isolates (TCH1516 and LAC). (A) Circular representation of whole genome comparison of *S. aureus* TCH1516 (internal ring) and LAC strains. Each ring of the circle represents a specific complete genome that corresponds to different colors in the legend on the right. The similarity between strains is represented by the intensity of the color. Darker colors represent higher similarities than lighter ones. Deleted regions are represented by blank spaces inside the circles. The whole genome comparisons were generated by BRIG. Alignment identity cutoffs of 0.8 (upper) and 0.5 (lower) were used to determine missing regions in the query genome (*S. aureus* LAC) compared to the *S. aureus* TCH1516 reference. Since *S. aureus* TCH1516 was the first genome to be annotated in the USA300 lineage, it was used as a reference genome for this study. (B) Dot plot of a nucleotide-based alignment of the genomes between *S. aureus* USA300 TCH1516 and LAC.

### Genome-wide identification of CodY-binding sites in *S. aureus* USA300 TCH1516

Previously, 57 CodY-binding sites in the *S. aureus* clinical strain UAMS-1 were identified *in vitro* deploying an affinity purification experiment using purified *S. aureus* His-tagged CodY and a related mutant strain (8). However, there are no direct *in vivo* measurements of the interaction between CodY and DNA in the recent USA300 isolates. Thus, using a novel monoclonal antibody, we performed Chromatin immunoprecipitation followed by exonuclease digestion (ChIP-exo) to identify the CodY-binding sites with single nucleotide resolution in *S. aureus* USA300 TCH1516 under RPMI with 10% LB medium (17).

Using a peak calling algorithm (MACE), a total of 165 CodY target genes (135 binding peaks) were identified from TCH1516 (**Figure 2A, upper panel**). As mentioned before, there were 57 CodY target genes identified in *S. aureus* UAMS-1 using IDAP-Seq, 57% (36/57) of which were also detected in TCH1516. Compared to the IDAP-Seq approach, this study detected the interaction between TF-DNA *in vivo* to enrich the DNA bound by CodY in its natural state, but also showed the location of each peak at single nucleotide resolution.

**Figure 2.**
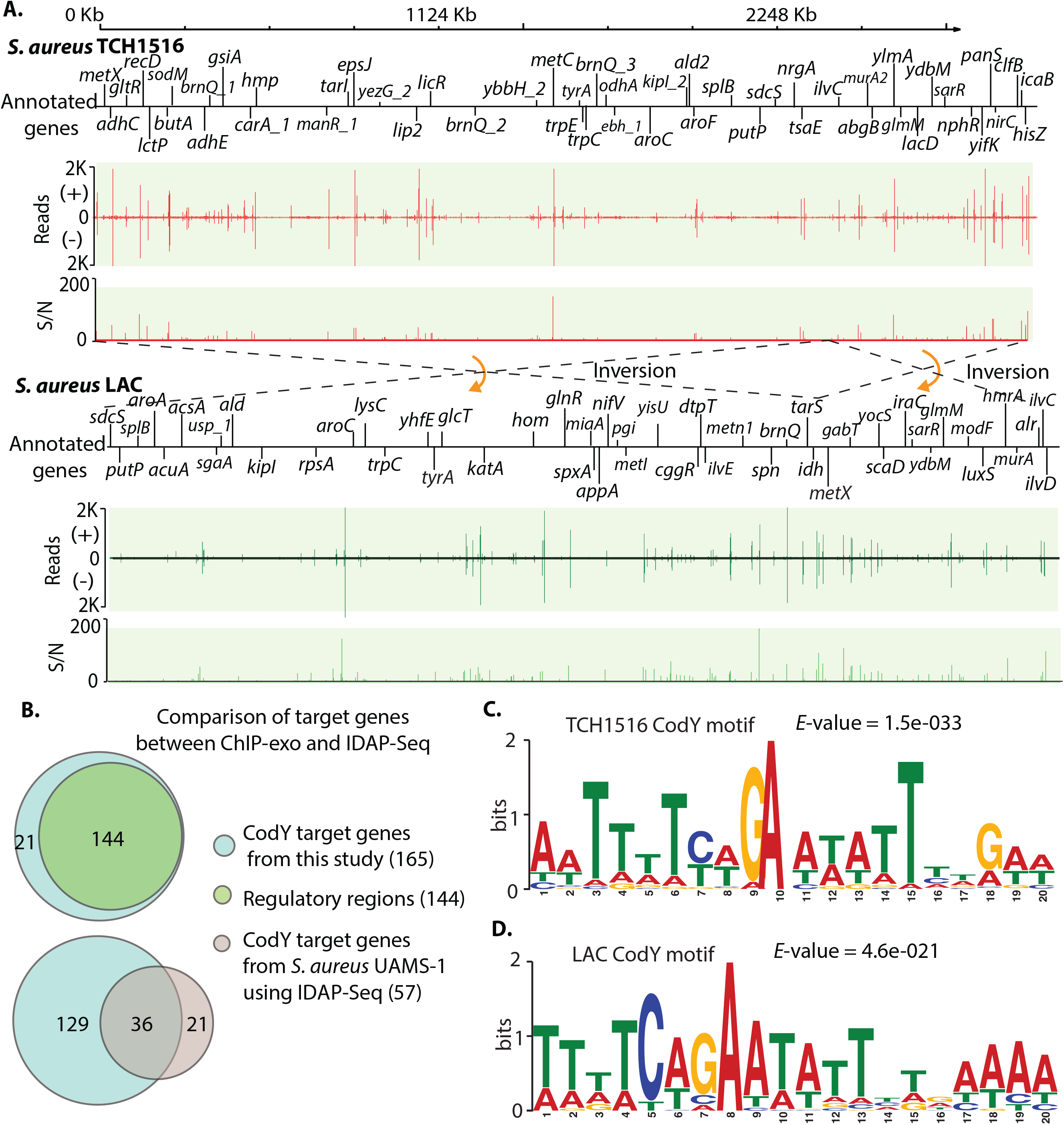
Comparison of CodY-binding sites between TCH1516 and LAC strains. (A) Direct comparison of CodY target genes between *S. aureus* substr. USA300 TCH1516 and LAC at the genome. Upper panel: an overview of CodY-binding profiles across the *S. aureus* TCH1516 genome at mid-exponential growth phase in RPMI 1640 + 10% LB medium. S/N denotes signal-to-noise ratio. (+) and (-) indicate forward and reverse reads, respectively. Bottom panel: an overview of CodY-binding profiles across the *S. aureus* LAC strain genome at mid-exponential growth phase under RPMI 1640 + 10% LB medium. (B) Distribution of *in vivo* CodY genome-wide binding sites at the genome of the TCH1516 strain (upper panel). Comparison of CodY-binding sites obtained from this study (ChIP-exo) with CodY target genes from *S. aureus* USA200 UAMS-1 using IDAP-Seq (bottom panel). (C) The consensus DNA sequence for *S. aureus* TCH1516 CodY binding motif. (D) The consensus DNA sequences for *S. aureus* LAC CodY binding motif.

Our results showed that 88% (144/165) of CodY binding sites were located within the intergenic regions, and the remaining 12% of binding sites were found in coding regions (**Figure 2B, upper panel**). Most of the binding sites located in intergenic regions were present upstream of assigned genes, indicating that CodY may play critical regulatory roles in the expression of these genes. A total of 128 novel CodY target genes were identified (**Figure 2B, bottom panel**). These findings expanded the list of CodY target genes in TCH1516 and enabled a better understanding of the global regulatory role of CodY in the USA300 lineage.

### Identification of the CodY-binding motif in *S. aureus* TCH1516

To identify the DNA sequence motif of CodY-binding sites, we used the MEME motif-searching algorithm with the genomic sequences of binding sites, and then identified the conserved 20-bp CodY binding motif, which was consistent with the previously characterized CodY DNA-binding consensus sequence (AATTTTCWGAAAATT) in *S. aureus* UAMS-1 (**Figure 2C**). Furthermore, the *S. aureus* TCH1516 CodY binding motif is similar with the CodY binding motif (AATTTTCWGAATATTCWGAAAATT) reported in *Listeria monocytogenes* and *Bacillus subtilis* (18, 19). These results suggest that CodY likely has a conserved DNA-binding domain in Gram-positive bacteria (**Supplementary Figure 1**).

### Comparison of *in vivo* CodY-binding sites in *S. aureus* TCH1516 and LAC strains

As two community-associated methicillin-resistant strains, *S. aureus* TCH1516 and LAC have an identical CodY sequence (**Supplementary Figure 2**). To investigate the direct gene targets of CodY, we employed the same monoclonal antibody to perform the ChIP-exo assay in *S. aureus* LAC under RPMI with 10% LB medium (17).

*S. aureus* LAC has 165 CodY target genes identified at the genome that were identical to those from the TCH1516 strain, though their positions at each chromosome are different due to the inversions mentioned earlier (**Figure 2A, bottom panel**). The alignment of the binding motifs revealed that the two strains have 18 nucleotides overlap (*p*-value= 9.35e-09) (**Supplementary Figure 3**). Among the 165 target genes, there are ten genes directly related to virulence in *S. aureus* TCH1516 and LAC strains (**Table 2**), consistent with the report that CodY links metabolism with virulence gene expression (6). Taken together, these data demonstrated that the global regulator CodY controls the expression of metabolism and virulence genes in *S. aureus* (9).

**Table 2.**
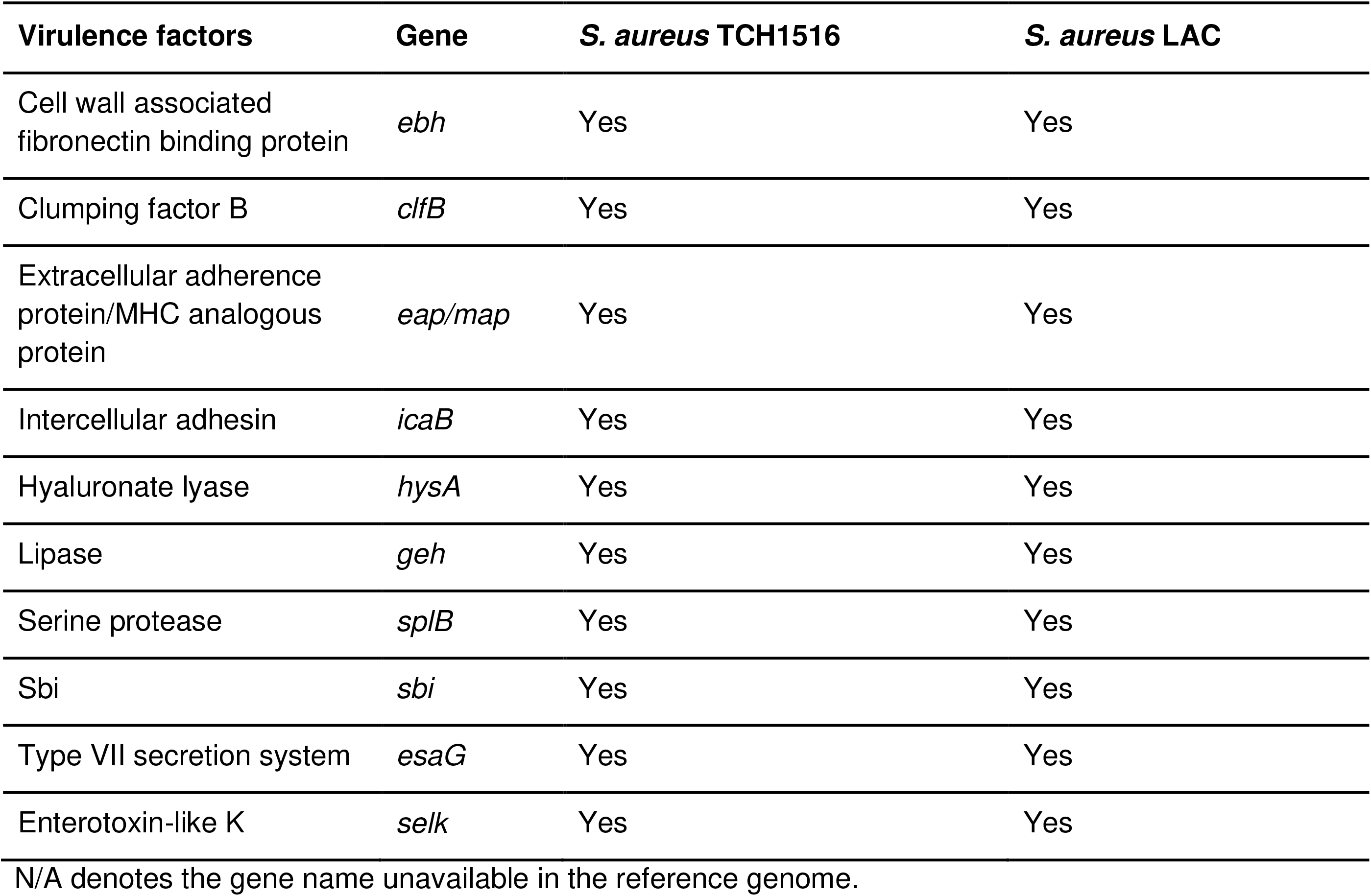
The CodY target genes considered to be virulence factors in the TCH1516 and LAC strains

### Strain-specific binding intensity is associated with DNA sequence variations in the binding site CodY

Though the genome-wide CodY binding sites identified were the same in the two strains, we found that strain-specific binding peaks had differing intensities (**Supplementary Figure 4**). The data showed that some of the binding peaks in TCH1516 had a higher intensity for CodY than those in LAC, and vice versa. 45 of 135 binding sites had identical DNA sequences (**Supplementary Figure 5**).

To fully evaluate whether the strain-specific binding peaks are due to the changes in the sequence-specific affinity to which CodY is bound, we first identified a 20-bp motif based on a merged set of CodY binding peaks from the strains using MEME (20) (**Supplementary figure 6**). We then utilized the computational method MAGGIE to measure differences in the DNA sequence in paired binding sites in the two strains (21). Among 135 pairs of peaks between the strains, 33.3% (45/135) had a range of motif score difference between −1 and 1. We plotted the pairs of peak heights for the 135 shared binding sites in LAC and TCH1516 (**Figure 3A**). When we highlighted the 45 binding sites with near identical DNA sequences, we observed that they represent the dots closest to the 45 degree line. This shows that these 45 sites have similar peak heights in the two strains. The binding sites where the DNA sequence differs are off the 45 degree line, showing that these differences lead to differential peak heights for the same binding site in the two strains.

**Figure 3.**
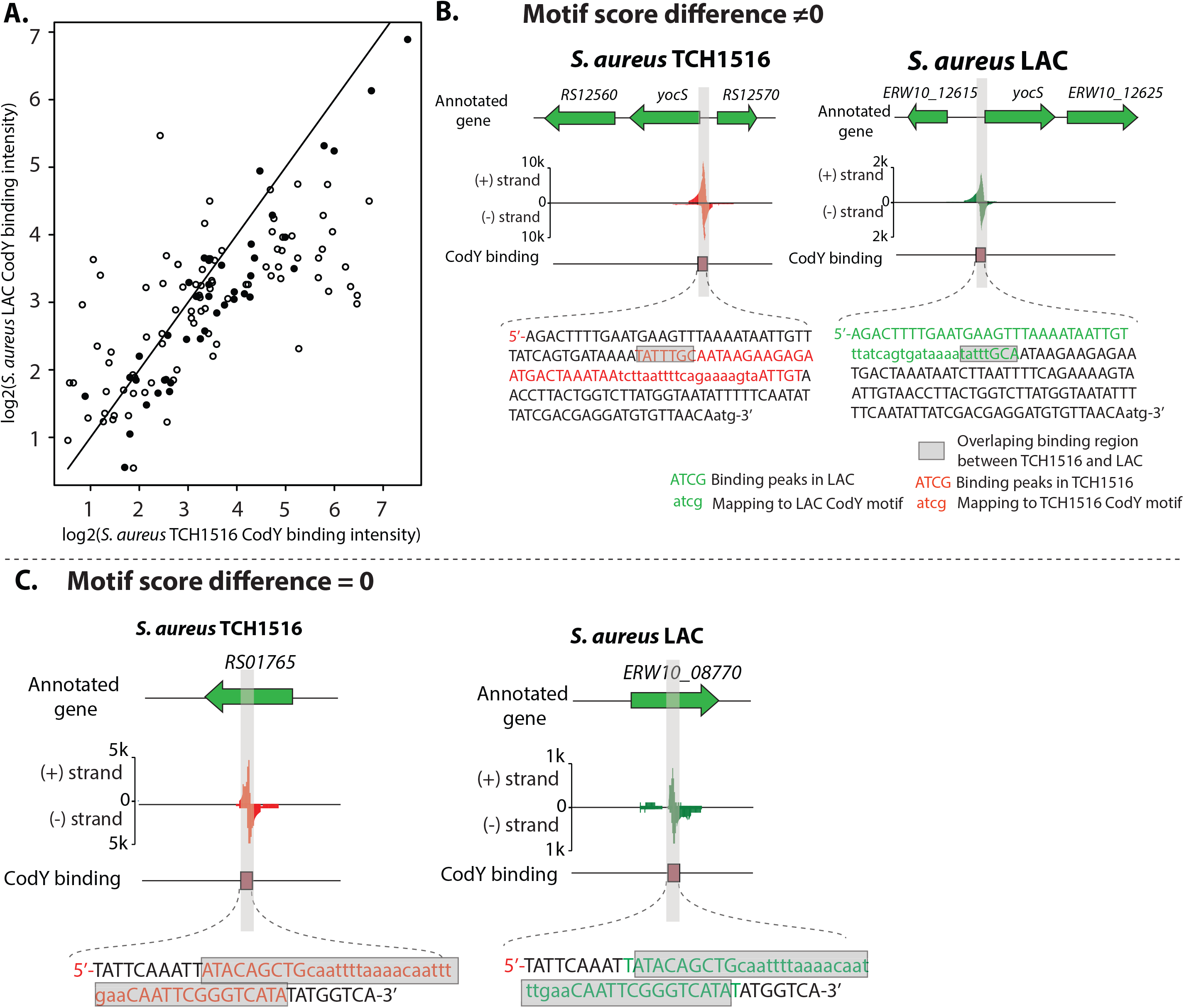
Differential CodY binding peaks area plot for TCH1516 and LAC highlighting binding sites with identical binding motifs. (A)Binding peak areas of log2(TCH1516_binding_intensity) (x-axis) and log2(Lac_binding_intensity) (y-axis) is shown. The diagonal line represents identical peak areas. The 45 binding sites with near zero sequence difference (Supplemental Figure 5) are shown with the solid dots, while those that are different are shown with open circles. (B)Case study I: the binding peak at the upstream of gene *yocS*, which has non-zero motif score difference between TCH1516 and LAC. Annotations of color code nucleotides are shown in the legend. Nucleotides in red and green represent the CodY peak sequences in TCH1516 and LAC, respectively. Grey denotes the overlap between a pair of peak sequences. (C) Case study II: the binding peak at the gene USA300HOU_RS01765, which has zero motif score difference between TCH1516 and LAC.

Further, to visualize the differences between a pair of binding peaks, we mapped CodY binding peaks to the reference genome. For example, the sequence of the peak located at the upstream of gene *yocS* in TCH1516 slightly (8 bp) overlaps the corresponding peak sequence in LAC **(Figure 3B)**. Another example is from a pair of peaks from unknown genes USA300HOU_RS01765 and ERW10_08770. They had zero motif score difference, and thus we observed that both peak sequences nearly overlap each other **(Figure 3C)**.

### Genome-wide reconstruction of CodY regulons in the *S. aureus* USA300 lineage

The ChIP-exo datasets from this study expanded the size of target genes to 165 in *S. aureus* USA300 lineage, which included 128 novel CodY target genes. Of these, 37% (47 of 128) were metabolic genes. Nearly all of these 47 metabolic genes were non-essential genes in *S. aureus* USA300.

To further characterize the regulatory roles of CodY in the *S. aureus* USA300 lineage, we compared gene expression profiling of the *codY* mutant to that of the wild type under RPMI with 10% LB medium, and found there were 809 genes differentially expressed in the *codY* mutant (at least 2-fold change (P<0.05) in RNA-seq expression) (**Figure 4A, Supplementary Figure 7**). The majority of genes related to metabolism were up-regulated in the *codY* mutant, which was consistent with the repressor role that CodY plays in *S. aureus* (22). Further, we used functional enrichment analysis by Clusters of Orthologous Groups (COG) classification of 809 differentially expressed genes to discover the top six differential enrichment pathways: amino acid transporter and metabolism, inorganic ion transport and metabolism, translation, ribosomal structure and biogenesis, transcription, carbohydrate transport and metabolism, and energy production and conversion (**Figure 4B**).

**Figure 4.**
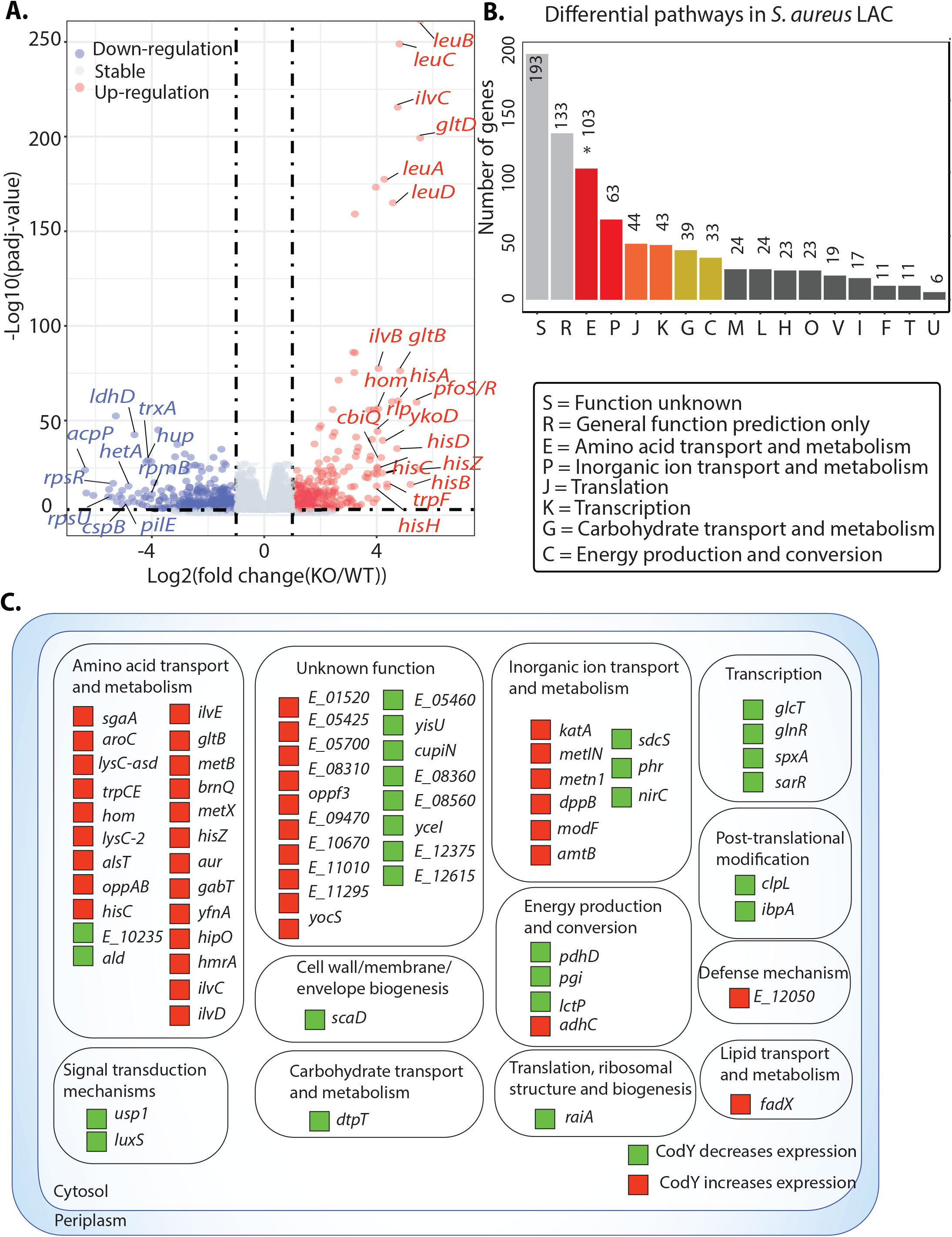
Role of CodY in regulation of *S. aureus* USA300 LAC genes differentially expressed in the *codY* mutant. (A) The *S. aureus* USA300 LAC direct CodY regulon. LAC genes that had a CodY ChIP-exo binding and had at least 2-fold change (P<0.05) in RNA-seq expression between the codY mutant to the wild-type strain were assigned to the direct regulon. (B) Functional enrichment analysis by Clusters of Orthologous Groups (COG) classification of 809 differentially expressed genes in *S. aureus* LAC *codY* mutant compared to wild type. The number of genes are based on the annotated genome. The top six enriched pathways were amino acid transport and metabolism, inorganic ion transport and metabolism, translation, ribosomal structure and biogenesis, transcription, carbohydrate transport and metabolism, and energy production and conversion. The functional enrichment was analyzed by performing the hypergeometric test. The asterisk indicates hypergeometric P-value <0.05. (C) Reconstruction of 72 CodY regulon in *S. aureus* LAC strain.

Next, to reconstruct the CodY regulon, we compared the expression profiling of the *codY* mutant to the wild type strain. Combining genome-wide target genes with transcriptomics, we expanded the size of CodY regulons to 72 target genes that were directly regulated by CodY (**Figure 4C**). Over half of the regulons (51%, 37 of 72) were related to the metabolic pathways (amino acid transport and metabolism, inorganic ion transport and metabolism, and energy production/conversion). In addition, 57% (41 of 72) of regulons were negatively regulated by CodY in the wild type. We also found that 90% (65/72) of CodY regulons had been identified in the CodY iModulon, which confirmed the regulatory roles of CodY (**Supplementary Figure 8**) (23). Further, CodY regulons were directly involved in signal transduction mechanisms, transcription, translation, post-translational modification, and defense mechanisms. These data indicated that CodY contributes to global regulatory roles beyond the metabolism of *S. aureus* USA300.

### Rerouting of flux through central carbon metabolism to generate the branched-chain amino acids (BCAAs)

CodY is reported to regulate the expression of metabolic genes in response to changes in the pools of specific metabolites, i.e., the branched-chain amino acids (BCAAs; isoleucine, leucine, and valine (ILV) and nucleoside triphosphate GTP), to regulate genes involved in the biosynthesis these amino acids (24). In addition to providing building blocks for proteins, BCAAs are also incorporated into the membrane as a part of branched chain fatty acids (BCFAs) (25). Therefore, sufficient uptake or biosynthesis of BCAAs is a key component of cellular homeostasis. Analysis of the codY iModulon and ChIP-exo data indicated that codY directly regulates genes required for both the transport and biosynthesis of BCAAs. For many other amino acids (e.g. histidine, aspartate, threonine), our data indicates that codY only directly regulates their biosynthetic genes and not their transport. Therefore, we sought to understand how *S. aureus* deals with BCAA starvation. In order to understand the partition of fluxes required for BCAA synthesis we utilized parsimonious Flux Balance Analysis (pFBA), using a previously published metabolic model of *S. aureus* USA300 (26, 27). pFBA determines metabolic fluxes that maximize growth while minimizing the sum of fluxes through the system (28). Simulated growth in RPMI supplemented with BCAA led to growth with zero flux through the BCAA biosynthetic pathway in favor of direct transporters of the necessary metabolites. However, restricting flux through all BCAA transporters (see Materials and Methods), led to a spike in flux through the biosynthetic enzymes (**Figure 5**). Interestingly, aspartate, a precursor to isoleucine biosynthesis, was not generated by the TCA cycle intermediate oxaloacetate. Instead, aspartate was derived from the breakdown of asparagine, while flux through the TCA cycle was redirected towards generating pyruvate via malate. This result indicated that in the presence of sufficient asparagine and aspartate in the media, *S. aureus* generates pyruvate to increase precursor pools available for isoleucine biosynthesis. Indeed, flux through the two enzymes linking the TCA cycle and aspartate, malate dehydrogenase, and aspartate transaminase, increased when aspartate and asparagine transporters were blocked in addition to BCAA biosynthesis.

**Figure 5.**
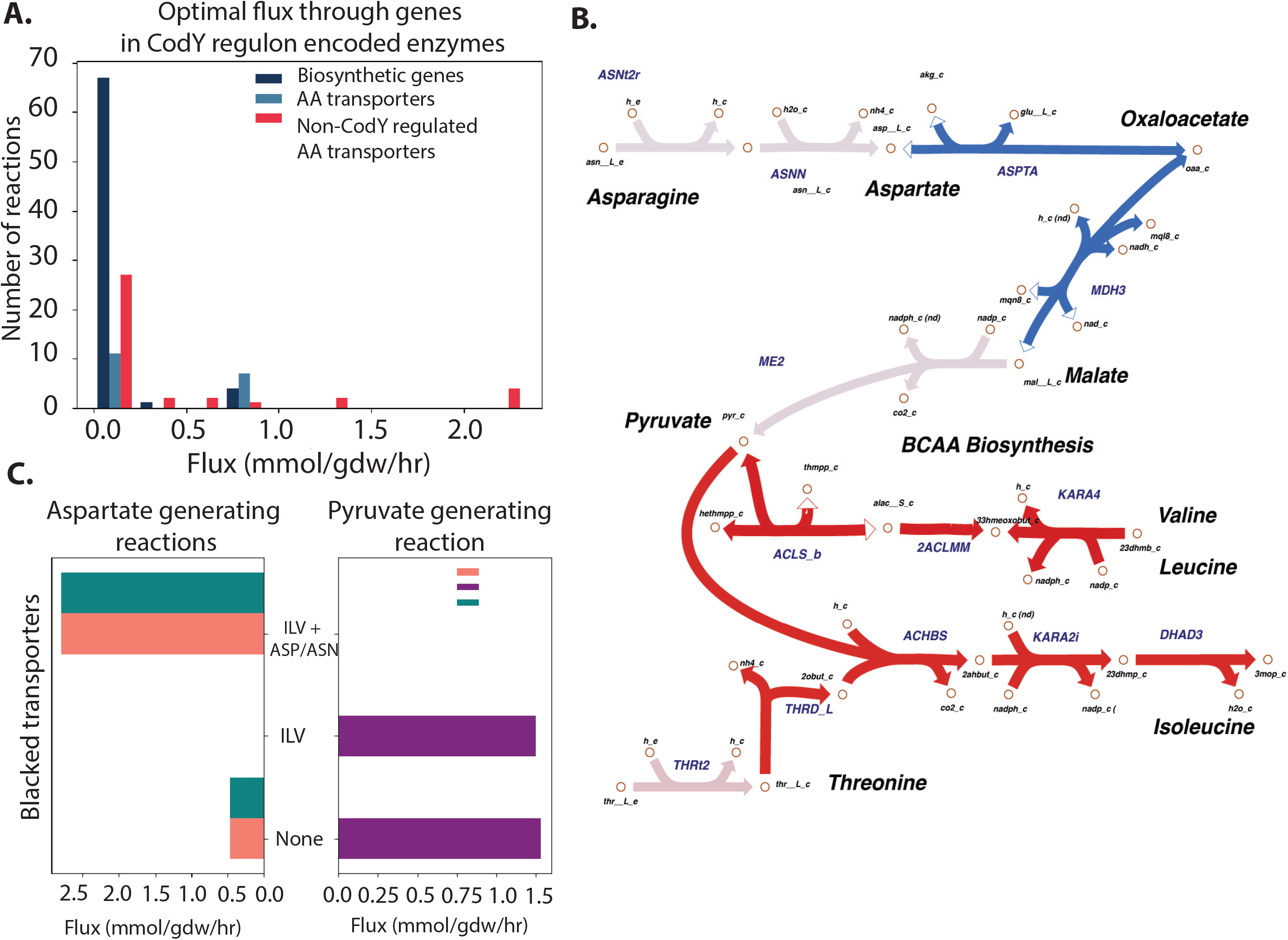
Central metabolic flux rerouting to generate BCAA. (A) Optimal flux through CodY regulated biosynthetic enzymes, transporters and non-CodY regulated transporters in RPMI medium. When amino acids are present, *S. aureus* imports them via transporters (light blue and red bars). (B) pFBA solution of *S. aureus* grown in a chemically defined medium (CDMG) without BCAA predicts rerouting of several central carbon metabolic fluxes to generate BCAA precursors for BCAA. Red arrows represent reactions with increased flux during BCAA starvation and blue arrows represent those with decreased flux relative to starvation conditions. Malate dehydrogenase and aspartate synthase (blue arrows) lower flux when BCAA transporters were blocked. Note: Full ILV biosynthesis pathways are not shown. (C) Flux through pyruvate generating malate enzyme (ME2) and aspartate generating malate dehydrogenase (MDH3) and aspartate transaminase (ASPTA). The pathways in bold correspond to the biosynthesis pathways that contain CodY target genes.

## Discussion

With the increasing number of whole genome sequences becoming available for multiple strains of a pathogenic species, the importance of the differences in their genomes and gene content is becoming more appreciated (29). Although many properties of pathogenic strains can now be predicted from sequence alone (30–32), detailed experimental characterization of differences for multiple strains is also needed. In this study, we combined genome-wide experiments and computational modeling to address the differences of CodY in the dominant CA-MRSA USA300 clinical isolates TCH1516 and LAC (15).

The study resulted in a series of significant findings. First, through the genome genome-wide identification of binding sites, we found the same 165 CodY target genes in both strains. Second, an examination of the differential binding intensity for the same target genes under the same conditions revealed that the variance was due to DNA sequence differences in the same CodY binding site in the two strains, while the *codY* protein was identical. This finding gives insights into the system-level analysis of CodY target genes and differential binding intensity across closely related strains. Third, the study identified ten virulence genes that belong to different types of virulence factors, which demonstrated that CodY connects metabolism genes with virulence genes in *S. aureus* (6). Considering that different *S. aureus* lineages may have distinct virulence factors, we could expand this study to identify many other virulence factor genes coordinated by CodY across different *S. aureus* strains. Finally, a genome-scale model of the metabolic network can be constrained by the regulatory action of CodY, and the results show the subsequent systematic rerouting of metabolism and pathway use.

Taken together, this study demonstrates, for the first time, the differences in the function of a conserved globally acting transcription factor (e.g., CodY) between closely related pathogenic strains. These results enable a wider range of studies to further decipher both subtle and major differences between the phenotypes of sequenced strains. This study motivates the characterization of common pathogenic strains and an evaluation of the possibility of developing specialized treatments for major strains circulating in the population.

## Materials and Methods

### Bacterial strains, plasmids, and culture conditions

The bacterial strains used in this study were described in Dataset 1. Methicillin-resistant *Staphylococcus aureus* (MRSA) strain substr. USA300 TCH1516 (also named USA300-HOU-MR) was originally isolated from an outbreak in Houston, Texas and caused severe invasive disease in adolescents (33). MRSA USA300 LAC was originally isolated from the Los Angeles county jail (15). *S. aureus* JE2, as a parental strain, was used for all sequence-defined Tn mutagenesis experiments. *CodY* mutant was from the Nebraska Transposon Mutant Library in which each of non-essential genes were disrupted via mariner Tn mutagenesis. All *S. aureus* strains were grown in tryptic soy broth (TSB, Sigma-Aldrich) or RPMI-1640 (Gibco) with 10% Lysogeny broth (LB, Sigma-Aldrich, MO) containing 10 g/L peptone, 5 g/L yeast extract, 10 g/L NaCl with the appropriate antibiotics for plasmid maintenance or selection (ampicillin, 100 ug/mL; chloramphenicol, 10 ug/mL) with shaking (250 rpm) at 37°C, maintaining a flask-to-medium volume ratio of 9:1, unless otherwise specified.

### Monoclonal antibody

The entire *S. aureus* CodY coding sequence was amplified, and introduced into *E. coli* BL21. The resulting glutathione-S-transferase (GST):: phoP fusion constructs were verified by DNA sequencing. To obtain the GST fusion proteins, *E. coli* cells were grown in 2 × YT medium at 30°C in an orbital shaker (180 rpm) to an OD_600_ of 0.6. Expression was induced with IPTG (0.1 mM final concentration) for 5 h. Cells were harvested by centrifugation, washed twice with PBS, lysed by sonication and then mixed with Glutathione Sepharose-4B beads (Pharmacia Biotech). Proteins were eluted with 10 mM reduced glutathione (in 50 mM Tris·HCl, pH 8.0) following the manufacturer’s recommendations and conserved in 40% glycerol at −80°C before use. Monoclonal anti-CodY antibody was generated by injection of CodY into the BALB/C mouse (Genscript, USA). Mouse anti-CodY IgG monoclonal antibody (IgG) was generated and purified (Genscript, USA).

### ChIP-exo experiments

ChIP-exo experimentation was performed following the procedures described previously (34). To identify CodY binding maps for each strain *in vivo*, the DNA bound to CodY from formaldehyde cross-linked cells were isolated by chromatin immunoprecipitation (ChIP) with the specific antibodies that specifically recognize CodY (Genscript, USA), and Dynabeads Pan Mouse IgG magnetic beads (Invitrogen) followed by stringent washings as described previously. ChIP materials (chromatin-beads) were used to perform on-bead enzymatic reactions of the ChIP-exo method. Briefly, the sheared DNA of chromatin-beads was repaired by the NEBNext End Repair Module (New England Biolabs) followed by the addition of a single dA overhang and ligation of the first adaptor (5’-phosphorylated) using dA-Tailing Module (New England Biolabs) and NEBNext Quick Ligation Module (New England Biolabs), respectively. Nick repair was performed by using PreCR Repair Mix (New England Biolabs). Lambda exonuclease-and RecJf exonuclease-treated chromatin was eluted from the beads and overnight incubation at 65 °C reversed the protein-DNA cross-link. RNAs- and Proteins-removed DNA samples were used to perform primer extension and second adaptor ligation with following modifications. The DNA samples incubated for primer extension as described previously were treated with dA-Tailing Module (New England Biolabs) and NEBNext Quick Ligation Module (New England Biolabs) for second adaptor ligation. The DNA sample purified by GeneRead Size Selection Kit (Qiagen) was enriched by polymerase chain reaction (PCR) using Phusion High-Fidelity DNA Polymerase (New England Biolabs). The amplified DNA samples were purified again by GeneRead Size Selection Kit (Qiagen) and quantified using Qubit dsDNA HS Assay Kit (Life Technologies). Quality of the DNA sample was checked by running Agilent High Sensitivity DNA Kit using Agilent 2100 Bioanalyzer (Agilent) before sequenced using HiSeq 2500 (Illumina) following the manufacturer’s instructions. Each modified step was also performed following the manufacturer’s instructions. ChIP-exo experiments were performed in biological duplicates.

### Peak calling for ChIP-exo dataset

Peak calling was performed as previously described (34). Sequence reads generated from ChIP-exo were mapped onto the reference genome using bowtie (35) with default options to generate SAM output files. MACE program was used to define peak candidates from biological duplicates for each experimental condition with sequence depth normalization (36). To reduce false-positive peaks, peaks with signal-to-noise (S/N) ratio less than 1.5 were removed. The noise level was set to the top 5% of signals at genomic positions because top 5% makes a background level in a plateau and top 5% intensities from each ChIP-exo replicates across conditions correlate well with the total number of reads (34, 37, 38). The calculation of S/N ratio resembles the way to calculate ChIP-chip peak intensity where IP signal was divided by Mock signal. Genome-scale data were visualized using MetaScope. (https://sites.google.com/view/systemskimlab/software?authuser=0)

### Motif search from ChIP-exo peaks

The sequence motif analysis for CodY binding sites was performed using the MEME software suite (20). For each strain, sequences in binding regions were extracted from the reference genome (*S. aureus* TCH1516: GenBank: NC_010079.1, NC_012417.1, and NC_010063.1; *S. aureus* LAC: GenBank: CP035369.1 and CP035370.1). To achieve a more accurate motif, the sequence of each binding site was extended by 10bp at each end. The width parameter was fixed at 20bp and the minsites parameter was fixed at 90% of the total number of the sequence. All other parameters followed the default setting.

### Clusters of Orthologous groups (COGs) enrichment

CodY regulons were categorized according to their annotated COG database (39). Functional groups in core, accessory, and unique CodY-regulated genes were determined by COG categories.

### Multiple genome comparison and alignment

MUMmer was used to run the complete nucleotide based alignments to check for synteny amongst the sequences (40). BLAST Ring Image Generator (BRIG) was used to show a genome wide visualization of coding sequences identity between *S. aureus* USA300 TCH1516 and LAC (41). Multiple genomes were analyzed by the M-GCAT, which is a tool for rapidly visualizing and aligning the most highly conserved regions in multiple prokaryote genomes. M-GCAT is based on a highly efficient approach to anchor-based multiple genome comparison using a compressed suffix graph (42).

### Determining Core Genome with Bidirectional BLAST Hits

To combine the data from the two strains, core genome-containing conserved genes between the LAC (GenBank: CP035369.1 and CP035370.1) and TCH1516 (GenBank: NC_010079.1, NC_012417.1, and NC_010063.1) were first established using bidirectional BLAST hits (43). In this analysis, all protein sequences of CDS from both genomes are BLASTed against each other twice with each genome acting as reference once. Two genes were considered conserved (and therefore part of the core genome) if 1) the two genes have the highest alignment percent to each other than to any other genes in the genome, and 2) the coverage is at least 80%.

### RNA Extraction and Library Preparation

*S. aureus* USA300 isolates JE2, and its derivatives of *codY* mutant were used for this study. The growth conditions and RNA preparation methods for data acquired from Choe et al. has been previously described(44). Detailed growth conditions, RNA extraction, and library preparation methods for other samples have also been already described (45). Briefly, an overnight culture of *S. aureus* was used to inoculate a preculture and were grown to mid-exponential growth phase (OD600 = 0.4) in RPMI+10%LB medium. Once in the mid-exponential phase, the preculture was used to inoculate the media containing appropriate supplementation or perturbations. All samples were collected in biological duplicates, originating from different overnight cultures. Samples for control conditions were collected for each set to account for batch effect.

### RNA-Seq Data Processing

The RNA-seq pipeline used to analyze and perform QC/QA has been described in detail previously (45). Briefly, the sequences were aligned to respective genomes, JE2, LAC or TCH1516, using Bowtie2 (46). The aligned sequences were assigned to ORFs using HTSeq-counts (47). Differential expression analysis was performed using DESeq2 with a P-value threshold of 0.05 and an absolute fold-change threshold of 2 (48). To create the final counts matrix, counts from conserved genes in LAC samples were represented by the corresponding ortholog in TCH1516. The counts for accessory genes were filled with 0s if the genes were not present in the strain (i.e., LAC-specific genes had counts of 0 in TCH1516 samples and vice versa). Finally, to reduce the effect of noise, genes with average counts per sample <10 were removed. The final counts matrix with 2,581 genes was used to calculate transcripts per million (TPM).

### Metabolic modeling and assessment of significant differences in flux distribution

We used BiGG model iYS854 and set the lower bound to the corresponding nutrient exchange to −1 mmol/gDW/h (the negative sign is a modeling convention to allow for the influx of nutrients) and −13 mmol/gDW/h for oxygen exchange (as measured experimentally) (27). Next, we compared two conditions with: 1) amino acid rich medium and 2) amino acid poor medium. In the first condition, assuming that in the presence of amino acid, CodY mediates the repression of multiple target genes. Specifically, we turned off the set of 65 reactions that CodY regulated related to biosynthetic pathways. No regulatory constraints were added. FBA was implemented with the biomass formation set as the functional network objective. Next, flux balance analysis was run in both conditions and the fluxes were sampled 10,000 times. All reaction fluxes were normalized by dividing by the growth rate to account for growth differences across the two media types. The flux distribution for each metabolic process was compared across both conditions using the Kolmogorov-Smirnov test, a non-parametric test which compares two continuous probability distributions. The distribution across two reactions was deemed to be significantly different when the Kolmogorov-Smirnoff statistic was larger than 0.99 with an adjusted p-value < 0.01.

### Motif mutation analysis

To gain insights into the CodY motifs from TCH1516 and LAC strains, MAGGIE was used for motif mutation analysis (21). For CodY, given a pair of CodY binding sequences of the same target gene from TCH1516 and LAC strains, the peak sequence with a higher binding intensity was considered as a positive sequence; the other one with a lower binding intensity was considered as a negative sequence. Here, CodY has 135 pairs of positive and negative sequences from TCH1516 and LAC. MAGGIE computes differences of representative motif scores (i.e., motif mutations) within each sequence pair by subtracting the maximal scores of negative from positive sequences and then statistically tests for the association between motif mutations and the differences in CodY binding intensity. Positive significant p-values from MAGGIE indicate that higher-affinity motif is associated with stronger CodY binding.

## Data availability

The ChIP-exo and RNA-seq datasets are accessible through GEO under accession number GSE159856 (review token: clgbaokcxhujhmt) and GSE163312 (review token: qfsvcukmnpwlvmb).

## Acknowledgements

We thank Richard Szubin for help with ChIP-exo library sequencing, Charles J. Norsigian for the insights about flux balance analysis of CodY, and Marc Abrams for reviewing and editing the manuscript.

## SUPPLEMENTARY MATERIALS

**Supplementary Figure 1** | **Comparison of CodY sequences among *S. aureus* USA300, *L. monocytogenes*, and *B. subtilis***. Red rectangle denotes the CodY DNA-binding domain (Helix-Turn-Helix domain).

**Supplementary Figure 2** | **Comparison of CodY sequences between *S. aureus* USA300 TCH1516 and LAC strains**.

**Supplementary Figure 3** | **The similarity between TCH1516 CodY binding motif and LAC CodY binding motif using TOMTOM. There are 18 bp nucleotides overlapping between them (*p*-value = 9**.**35e-09)**.

**Supplementary Figure 4** | **Comparison of CodY binding intensity between TCH1516 and LAC strains**.

**Supplementary Figure 5** | **Distribution of CodY peaks based on the range of motif score difference**. X-axis represents the range of motif score difference. Y-axis represents the number of peaks that fall into each range. Black-color bar showed 45 peaks with the motif score difference at the range of (−1, 1).

**Supplementary Figure 6** | ***S. aureus* USA300 CodY binding motif from a merged set of TCH1516 and LAC binding peaks**.

**Supplementary figure 7** | **A heatmap for wild type and *codY* mutant samples for expression profiling data from *S. aureus* USA300 LAC**.

**Supplementary Figure 8** | **Comparison of genes in the CodY regulon (red) and CodY iModulon (green). The number in the circle represents the amount of genes in the category**. CodY regulon is defined by a binding peak in the genes promoter region and detection of differential gene expression of the wild type and the *codY* mutant during growth in RPMI 1640 and 10% LB medium.

**Dataset S1 The strains used in this study**

